# Distinct dimensions of emotion in the human brain and their representation on the cortical surface

**DOI:** 10.1101/464636

**Authors:** Naoko Koide-Majima, Tomoya Nakai, Shinji Nishimoto

**Author notes:** Corresponding author: Shinji Nishimoto, Center for Information and Neural Networks, National Institute of Information and Communications Technology, Yamadaoka 1-4, Suita, Osaka 565-0871, Japan, E-mail address.

## Abstract

We experience a rich variety of emotions in daily life. While previous emotion studies focused on only a few predefined, restricted emotional states, a recent psychological study found a rich emotional representation in humans using a large set of diverse human-behavioural data. However, no representation of emotional states in the brain using emotion labels has been established on such a scale. To examine that, we used functional MRI to measure blood-oxygen-level-dependent (BOLD) responses when human subjects watched 3-h emotion-inducing movies labelled with 10,800 ratings regarding each of 80 emotion categories. By quantifying canonical correlations between BOLD responses and emotion ratings for the movie scenes, we found 25 significant dimensions of emotion representation in the brain. Then, we constructed a semantic space of the emotion representation and mapped the emotion categories on the cortical surface. We found that the emotion categories were smoothly represented from unimodal to transmodal regions on the cortical surface. This paper presents a cortical representation of a rich variety of emotion categories, which covers most of the emotional states suggested in traditional theories.

## Introduction

A central topic in affective neuroscience is to clarify how emotions are represented in the human brain. Recent functional magnetic resonance imaging (fMRI) studies have addressed this issue by showing brain representations of specific emotional states ^1–8^. However, the results were not sufficient to establish a brain representation of all of the emotional states that we experience because the emotional states were confined to those defined in the traditional emotion theories.

Traditionally, two main theories regarding the constitution of emotion have been postulated: One is the basic (categorical) emotion theory, which posits that emotional states can be explained by distinct categories of a few to 15 basic emotions (e.g. ‘fear’, ‘sadness’, ‘happiness’) ^9–13^. These categories have often been used in a hierarchical structure (e.g. featuring ‘anger’-related subcategories of emotions such as ‘annoyance’ and ‘fury’). However, when using this structure, it is difficult to represent the fuzzy boundaries among such emotion families ^14^. The other theory is the affective-dimension theory, in which emotion is explained in a continuous space consisting of a few dimensions (e.g. ‘arousal’ and ‘valence’) ^14–17^. However, such a low-dimensional model is inadequate to account for differences among multiple emotional categories such as anger and fear ^18^.

A recent study addressed the problems regarding the discriminability of the traditional emotion theories and provided a more natural interpretation of emotion by constructing a semantic space of emotion from reports of emotional experiences ^19^. The authors collected emotion categories, affective dimensions and free affective words for each of 2,185 movie clips. By examining the correlations and predictability between the rating types, the authors found 27 independent dimensions of specific emotion categories. The number of emotion dimensions, 27, is a richer variety of emotional states than in the traditional emotion theories. This behavioural observation suggests that the use of movie clips would enable us to measure brain activity associated with a rich variety of emotions.

To provide a comprehensive understanding of the brain representation of emotion, we used fMRI to measure blood-oxygen-level-dependent (BOLD) responses from eight subjects while they watched 3-h movies consisting of 720 clips that were selected to induce various types of emotion. The movies were rated regarding each of 80 emotion categories (see Methods), which constitutes greater variety than used in the previous fMRI studies ^1–8^. Using the emotion ratings, first, we examined the number of significant dimensions having high correlations between emotion ratings and the BOLD responses (see Methods). For each voxel, we estimated the BOLD responses to each of the 80 emotion categories by using a regularised linear regression. We constructed a semantic space consisting of dimensions of response patterns to represent the 80 emotion categories. Then, we performed a factor rotation analysis to give the interpretation of the dimensions and showed the cortical gradient of each voxel’s factor loadings.

**Figure 1.**
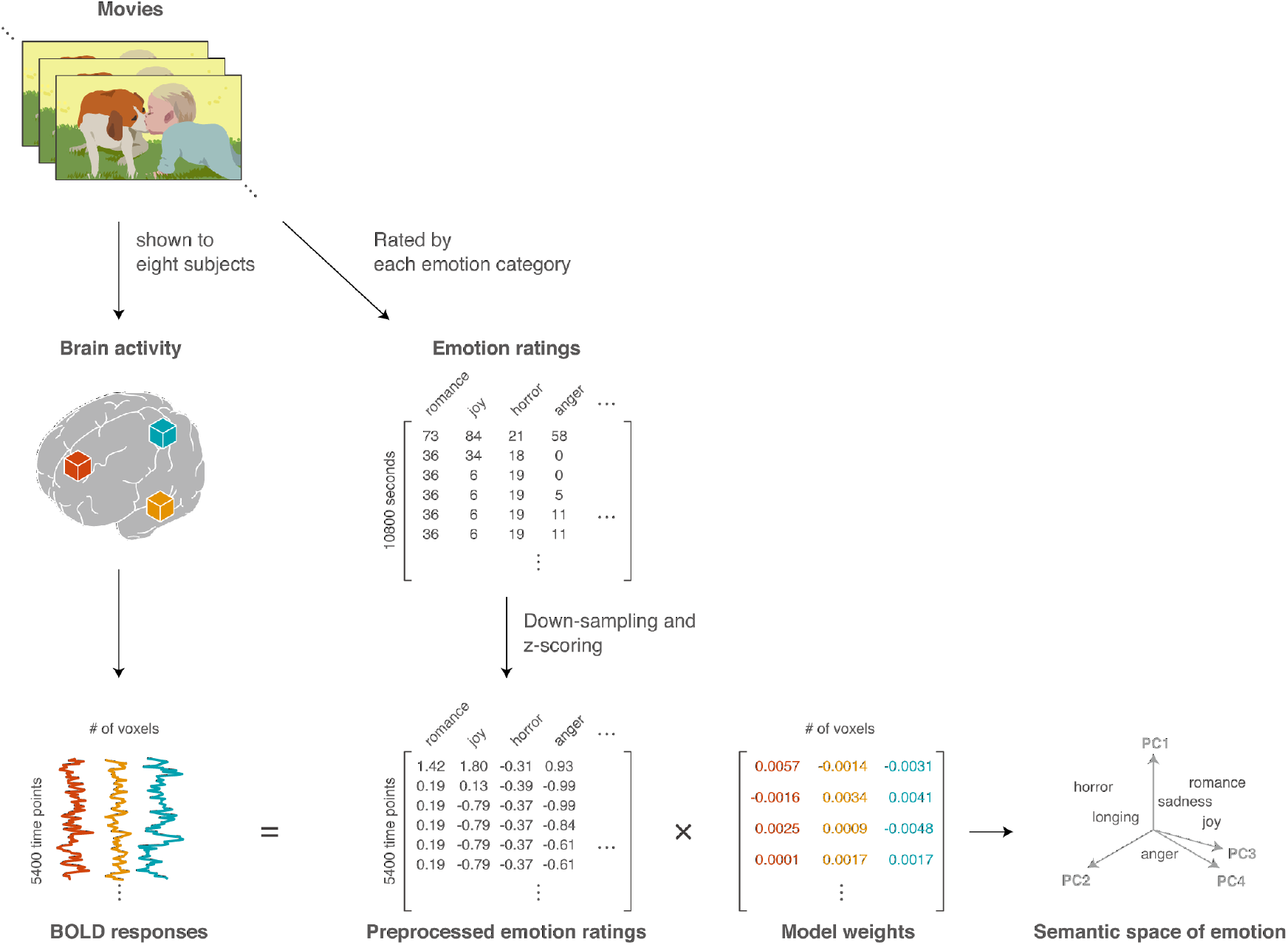
Schematic of experiment and procedure for constructing a semantic space of emotion. BOLD responses for eight subjects were measured while they watched emotion-inducing movies for 3 h. Each movie scene was rated regarding 80 emotion categories. Voxel-wise response was modelled as a linear weighted sum of the emotion ratings using an L2-regularised regression procedure. A semantic space was constructed by performing a dimension reduction on the estimated model weights.

## Result

### Twenty-five dimensions of emotion ratings were significantly correlated with the BOLD responses

First, we revealed significant dimensions of emotion ratings that were correlated with the BOLD responses. For this purpose, we used a canonical correlation analysis (CCA) between the ratings (3,600 time points × 80 dimensions) and the BOLD responses (3,600 time points × 3,984–13,068 voxels per subject) of the training dataset from each of the eight subjects (S1–S8), where factor loadings for each of the ratings and the responses were estimated to have the maximum correlation between them (see Methods). Then, we tested whether factor loadings showed a significant correlation (*p* < 0.01, with Bonferroni correction for 80 emotion categories) for each dimension of the ratings, using a test dataset of the ratings (1,800 time points × 80 dimensions) and the responses (1,800 time points × 3,984–13,068 voxels per subject). The results revealed 19~32 significant dimensions across subjects (S1: 20; S2: 27; S3: 25; S4: 28; S5: 23; S6: 32; S7: 24; S8: 19). The median was 24.75. We employed 25 dimensions in the next analysis to construct a semantic space of emotion.

### A semantic space shows brain representation of 80 emotion categories

To construct the semantic space of emotion, we used a dimensionality reduction technique on the BOLD-response patterns to 80 emotion categories for all the subjects. First, we concatenated estimated weights from individual subjects (80 × 18,684–34,066 voxels per subject). Then, we performed a principal component analysis (PCA) on the concatenated weights while treating the emotion categories and voxels as dimensions and samples, and reduced the dimensionality from the original 80 to 25. The semantic space of emotion was defined as the space consisting of the 25 dimensions. To maintain the quality of the semantic space, we only used voxels with high prediction performance of the regression model (see Methods).

In the semantic space of emotion, the distance for each pair of emotions represents the dissimilarity in the BOLD-response patterns between them. Fig. 2a shows the semantic space projected into the two-dimensional space, maintaining the emotion-pair distances in the 25-dimensional space as much as possible. In this space, positive and negative emotions are separated. As supporting results, we labelled each emotion in the semantic space as a positive, negative, or ambiguous emotion based on the emotion-word hierarchy of WordNet-Affect ^20^ (Fig. S1a). There, intra-class emotion labels are located close to each other. This tendency is quantitatively confirmed by using a random permutation test (Fig. S1b).

**Figure 2.**
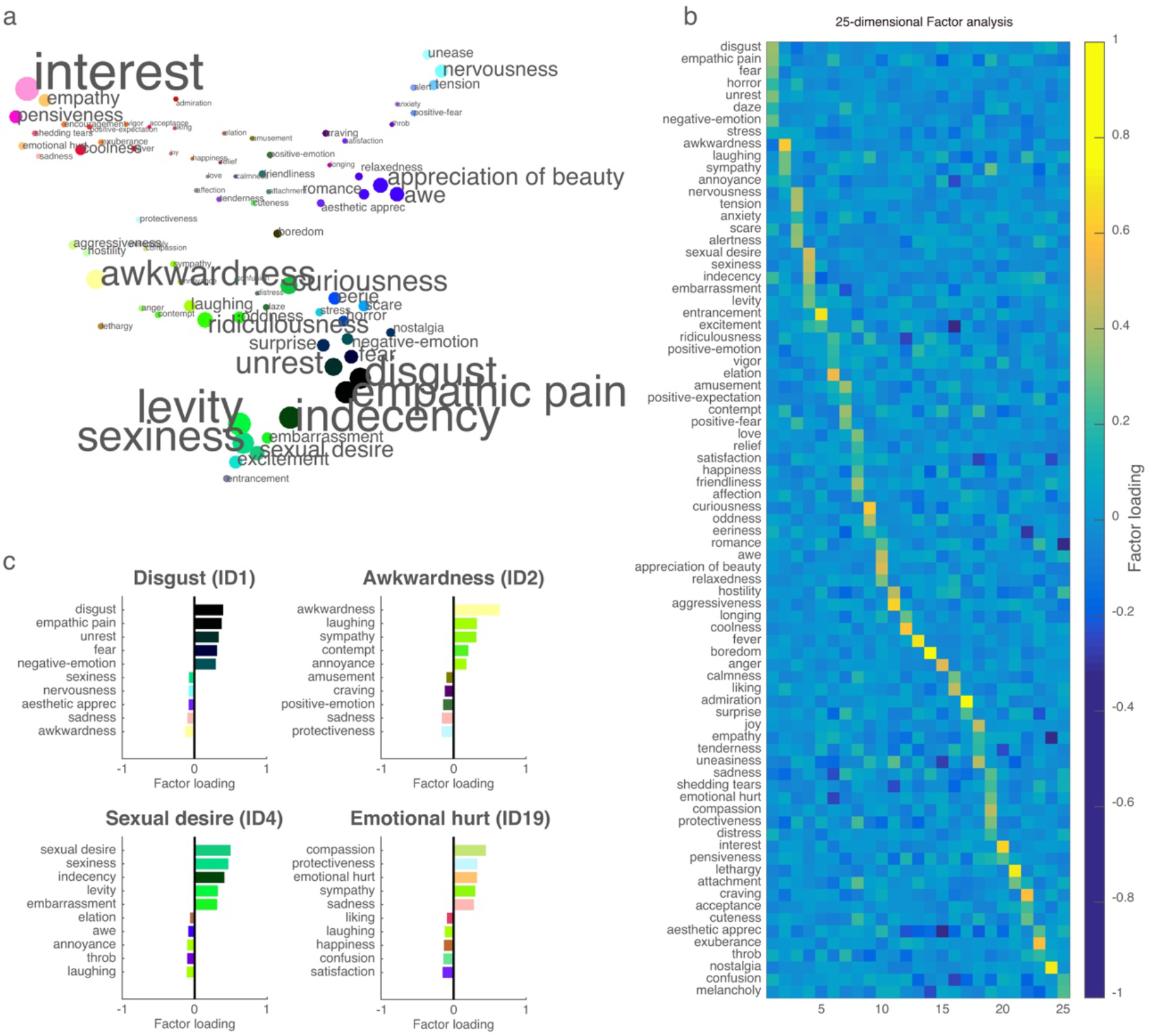
A semantic space of emotion and varimax factor loading. **a** Eighty emotion categories were plotted according to distances between the estimated weights of the L2-regularised regression model, using *t*-distributed stochastic neighbour embedding (*t*-SNE ^21^). A single RGB colour was assigned to each emotion category according to the three main components of the estimated weights (see Methods). Both marker size and font size were modulated according to the average weight across voxels (larger size denotes higher average weights). **b** Factor loadings of 25 components (= 25 emotion dimensions) of the estimated weights after varimax rotation. **c** Factor loadings of four representative components: ‘Disgust’, ‘Awkwardness’, ‘Sexual desire’ and ‘Emotional hurt’. The bars indicate the five highest and lowest factors.

To interpret each principal component of the semantic space, we performed varimax factor rotation (Fig. 2b). The results showed that each factor had high weights for related emotion categories such as ‘fear’ and ‘horror’. Fig. 2c shows factor loadings for the four representative factors, where the top-left factor (ID1) contains high weights in ‘Disgust-related’ emotions such as ‘disgust’, ‘empathic pain’ and ‘fear’. The top-right factor (ID2) contains high weights in ‘Awkwardness-related’ emotions such as ‘awkwardness’ and ‘laughing’. The bottom-left factor (ID4) contains high weights in ‘Sexual-desire-related’ emotions such as ‘sexual-desire’, ‘sexiness’ and ‘levity’. The bottom-right factor (ID19) contains high weights in ‘Emotional-hurt-related’ emotion categories such as ‘compassion’, ‘emotional hurt’ and ‘sadness’. Interpretations of the other factors are shown in Fig. S2. We found that each dimension was related to a specific emotion category, and more than half of the factors corresponded to emotion dimensions addressed in the work of Cowen et al. (2017): ‘Awkwardness’, ‘Sexual desire’, ‘Entrancement’, ‘Amusement’, ‘Adoration (Friendliness)’, ‘Boredom’, ‘Anger’, ‘Admiration’, ‘Joy’, ‘Emotional hurt (Sadness)’, ‘Interest’, ‘Craving’ and ‘Nostalgia’. The other dimensions also allowed a similar interpretation as those in the work of Cowen et al. (2017), although the grouping was different: In our case, negative emotions such as ‘disgust’, ‘empathic pain’ and ‘fear’ were combined into one dimension, ‘Disgust’. Furthermore, ‘tension’, ‘nervousness’, ‘scare’ and ‘positive-fear’ were combined into one dimension, ‘Nervous scared’, while distinct dimensions were reported for the related emotions (‘Anxiety’, ‘Horror’ and ‘Fear’) in the work of Cowen et al. (2017). As for positive emotions, we found some ‘Excitement-related’ emotions (‘excitement’, ‘entrancement’, ‘exuberance’, ‘encouragement’, and ‘fever’) that showed low contributions to explaining the BOLD responses (Fig. S3).

The semantic space was constructed from the aggregated regression model weights across all the subjects. To examine whether the obtained semantic space was consistent across individual subjects, we computed the Pearson’s correlation coefficient between the semantic space from one subject and that from the remaining seven subjects. As a control condition, we also computed the correlation coefficient between the semantic space from each excluded subject and that from the emotion ratings (see Methods). Table S1 lists the two types of correlation coefficients. For all of the subjects, we found a higher correlation between the individual and the group semantic space, than in the comparison with the emotion ratings. This suggests that the individual semantic space was consistent across the subjects and that the group semantic space can be used as a representative space for all the subjects.

### Cortical gradient of 25 emotion dimensions

To examine how emotion dimensions were represented on the cortical surface, first, we assigned an RGB colour according to the first to third principal components of the semantic space (see Methods). Two examples (subjects S1 and S5) of the visualisation are shown in Fig. 3a. (The cortical maps for the other subjects are shown in Fig. S4.) In both of the cortical maps, we found a weight gradient shown as a color gradient in the whole cortical surface. For example, different colours were assigned to the postcentral area and the superior temporal area. Indeed, weight distributions of 80 emotion categories differed between the two representative voxels obtained from these two areas (Fig. 3b). Specifically, one voxel from the postcentral sulcus contained high weights in ‘Disgust’-related emotions such as ‘empathic pain’ and ‘unrest’. The other voxel in the superior temporal gyrus contained high weights in ‘Interest’ and ‘Awkwardness’-related emotions such as ‘interest’, ‘empathy’, ‘laughing’ and ‘awkwardness’. This suggests that weight variability across voxels was successfully visualised across the cortical map.

**Figure 3.**
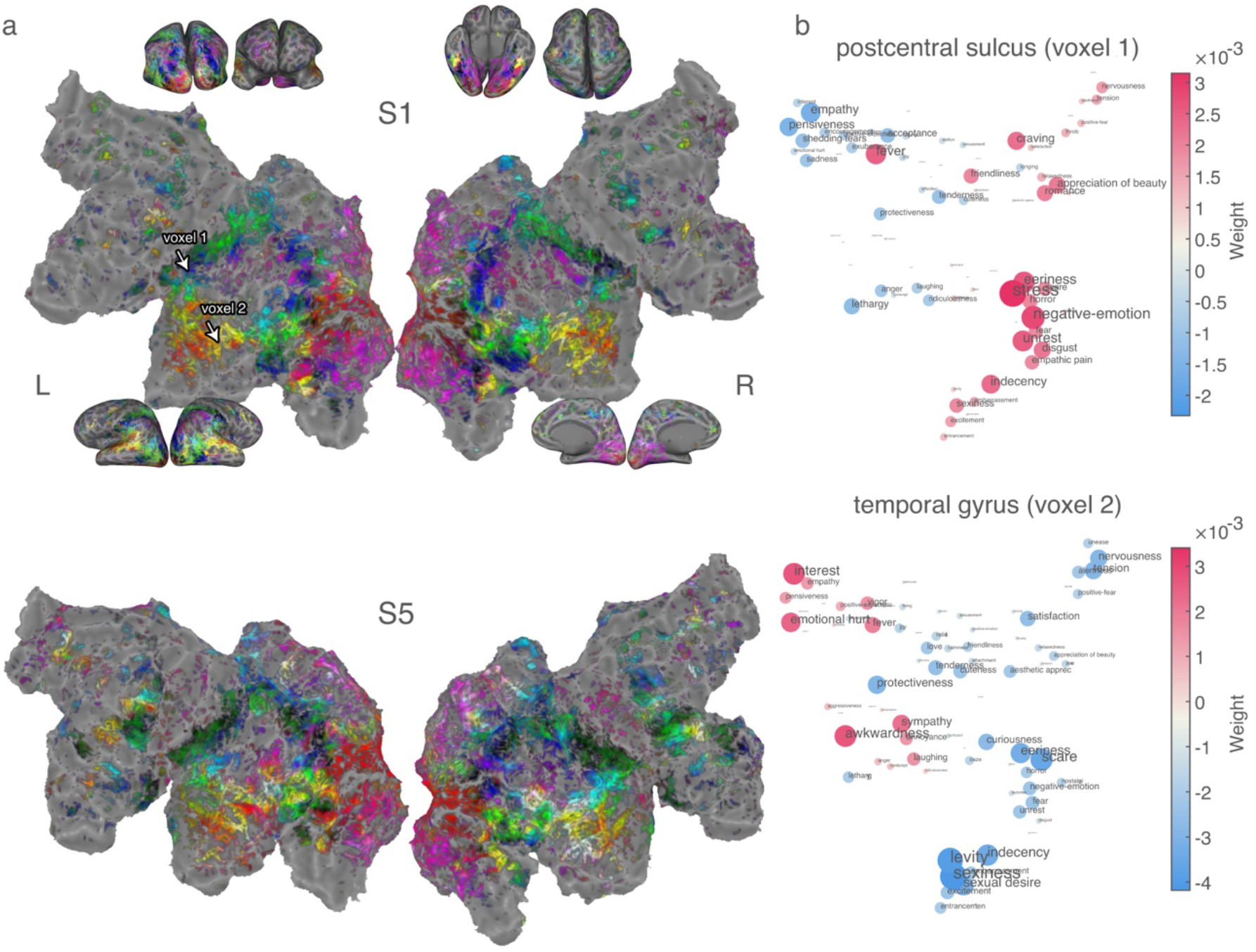
Cortical map of the semantic space of emotion. **a** Cortical maps for two subjects (S1 and S5). Emotion representations were visualised by assigning RGB colours according to the three main components of the semantic space (see Methods). In each cortical map, we show the results only for voxels showing significantly high prediction accuracy (p<0.0001, uncorrected). **b** Examples of the weight distribution for two voxels in the postcentral sulcus and the temporal gyrus of a single subject (S1). Positions of the two voxels are indicated on the S1’s cortical map in Fig. 3a. A red colour denotes positive weight and a blue colour denotes negative weight. Positions of emotion categories were the same as the positions in Fig. 2a.

Then, we examined how weights of the emotion dimensions varied across the cortical surface, especially in the four areas that showed consistent colour gradients across subjects, namely, the postcentral area, the superior temporal area, the inferior parietal area and the precuneus (Fig. 4b). In each of these four areas, we obtained weights of the 25 emotion dimensions from 30 successive positions on eight manually defined lines (line 1–8). The cortical maps of two subjects (S1 and S5) in Fig. 4a show the lines. The obtained weights from each line were plotted in a sequential order of spatial coordinates in the flattened map (Fig. 4b, Fig. S5). The postcentral area showed high weights in negative emotions such as ‘Disgust’ and ‘Nervous scared’ across the successive positions. ‘Sexual desire’ also showed high weight, and high correlation in the weight gradient with ‘Disgust’ (line1: *r* = 0.428, line2: *r* = 0.648). The superior temporal area showed a high weight gradient from ‘Interest’ to ‘Awkwardness’ and ‘Aggressiveness’. ‘Aggressiveness’ showed similar weight gradients from ‘Nervous scared’ (line3: *r* = 0.95, line4: *r* = 0.97), ‘Sexual desire’ (line3: *r* = 0.77, line4: *r* = 0.88) and ‘Curiousness’ (line3: *r* = 0.90, line4: *r* = 0.95). The inferior parietal area showed a high weight gradient from ‘Disgust’ to ‘Awe’ and ‘Interest’. ‘Disgust’ showed similar weight gradient from ‘Sexual desire’ (line5: *r* = 0.79, line6: *r* = 0.75) and ‘Curiousness’ (line5: *r* = 0.95, line6: *r* = 0.90). ‘Interest’ showed similar weight gradients from ‘Emotional hurt’ (line5: *r* = 0.93, line6: *r* = 0.77) and ‘Coolness’ (line5: *r* = 0.95, line6: *r* = 0.89). The precuneus showed a high weight gradient from ‘Interest’ and ‘Awkwardness’ to ‘Nervous scared’. ‘Interest’ showed a similar weight gradient to ‘Emotional hurt’ (line7: *r* = 0.92, line8: *r* = 0.80). Furthermore, ‘Sexual desire’ and ‘Disgust’ also showed high weights across the latter positions, and their gradients were highly correlated (line5: *r* = 0.94, line6: *r* = 0.96). As supporting results, we showed that each line was obtained from anatomically similar regions, which is supported by the same or neighbour labels of the Destrieux atlas ^22^ obtained from each line across all the subjects (Fig. S6).

**Figure 4.**
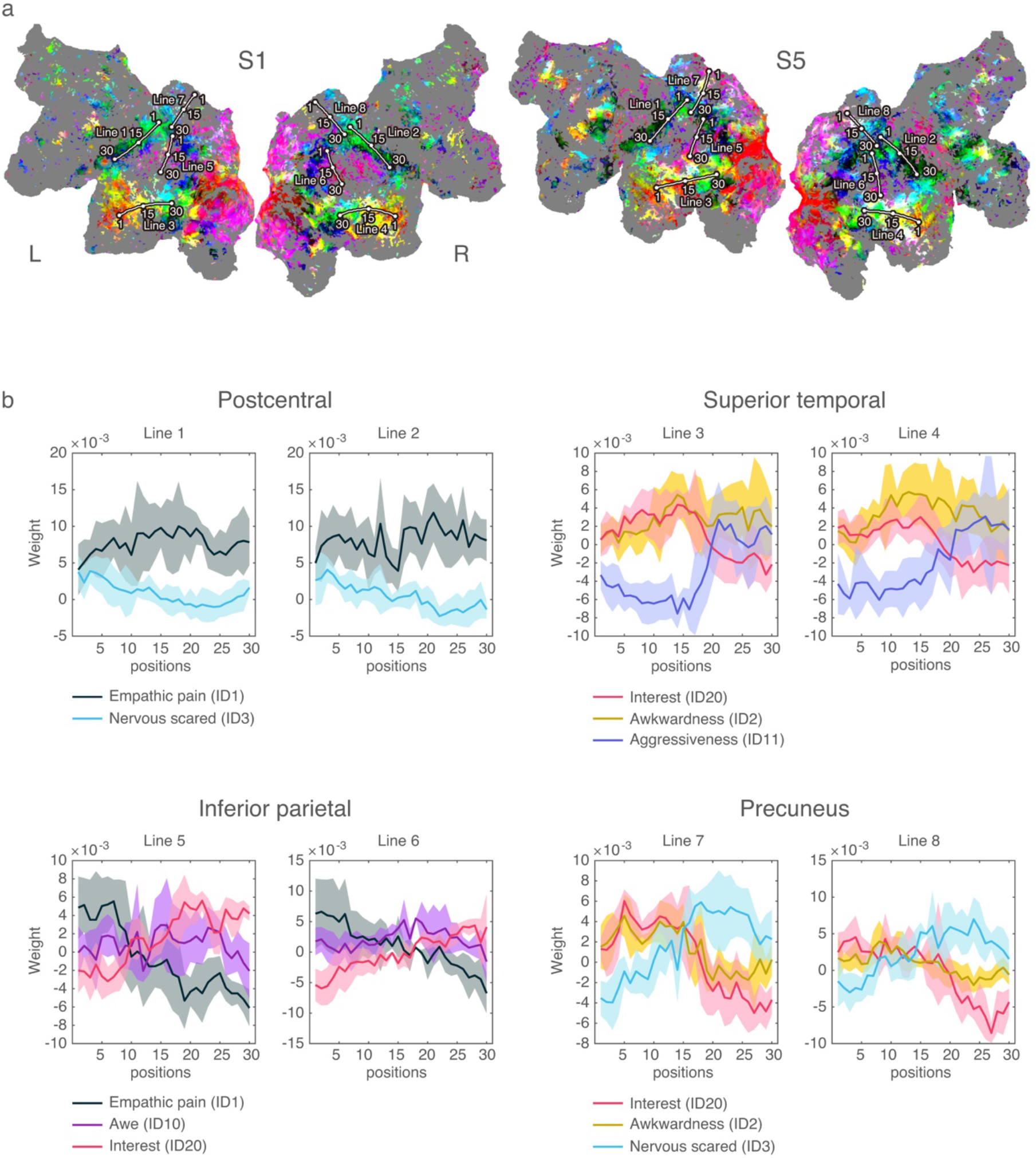
Weight gradient on the flattened cortical surface. **a** Positions of eight representative gradients are depicted as lines for two subjects (S1 and S5). Lines 1 and 2 were mainly located in the postcentral area, Lines 3 and 4 were mainly located in the superior temporal area, Lines 5 and 6 were mainly located in the inferior parietal area, and Lines 7 and 8 were mainly located in the precuneus. The anatomical location of each line was consistent across subjects, which is supported by results in Fig. S4. **b** Spatial transitions of representative (high absolute weights) emotion dimensions are plotted for each line. A bold plot denotes mean contribution across subjects. Shaded areas denote the standard deviation of contributions across subjects. The weights of all the emotion dimensions are shown in Fig. S5.

## Discussion

To reveal how a rich variety of emotion categories are represented in the human brain, we estimated the BOLD responses to 80 emotion categories, and constructed a semantic space consisting of dimensions of response patterns to specific emotion categories. We found 25 semantic dimensions that exhibited statistically significant signals. This is a considerably richer variety of emotions than the number of emotion categories or affective dimensions used in previous brain imaging studies (i.e. a few to 15) ^1–8^. Furthermore, each dimension showed a smooth gradient of the weights in the cortical surface. Such a gradient could not be revealed in these previous studies using a few emotion categories. By using a rich variety of emotions, we were able to construct a continuous space of emotion, and show more detailed localisation of the emotion categories.

The semantic space found here covered a large variety of emotion representations, and traditional emotion theories can be explained by part of the representations. In our semantic space, positive-emotion and negative-emotion categories were separately distributed (Fig. S1). This indicates that the positive/negative distinction is a basic factor for the organisation of emotions in the brain representation. This organisation is not contradictory to the emotion distribution in ‘Valence’ dimension of the core affect model ^14–17^. Furthermore, we found that most of the dimensions in our semantic space consisted of semantically related emotion categories that allowed interpretation as a group (e.g. ‘tension’, ‘nervousness’ and ‘alertness’ to form the ‘Nervous scared’ dimension). The total of 25 dimensions covered most of the emotion categories posited in the basic emotion theories ^9–13^. Some dimensions for negative emotions such as ‘Disgust’ and ‘Nervous scared (Fear)’ had higher average weights than those for the other dimensions such as ‘Joy (Happiness)’ and ‘Anger’ (Fig. S3). It has been suggested that negative emotions are more informative in the brain representation than the other categories, which may be because of their importance for survival^23, 24, 31^.

Emotion representation in our semantic space was partly consistent with the representation revealed using behavioural data. Specifically, the emotion-category distribution in our semantic space was consistent with the manually defined hierarchy of WordNet-Affect ^20^ (Fig. S1) in that the positive, negative and ambiguous emotions were distinguished from each other ^25^. This distinct positive/negative representation was also observed in a recent study using large-scale psychological assessments ^19^. Quantitatively, regarding the number of emotion dimensions, the 25 dimensions found here were similar to the 27 dimensions reported in the work of Cowen et al. (2017). Qualitatively, most of the 25 dimensions showed similar meanings to the behaviourally-defined emotion dimensions (e.g. ‘Awkwardness’, ‘Sexual desire’, and ‘Entrancement’). However, we found some differences regarding the boundaries among negative emotions (‘Empathic pain’, ‘Disgust’, and ‘Fear’). In our case, these negative emotions were combined into a single dimension while they were maintained as separate dimensions in the work of Cowen et al. (2017).

Regarding the emotion dimensions of the semantic space, we found the smooth gradients of the weights in the following areas: the superior temporal area, the inferior parietal area, the precuneus, and the postcentral area. A previous study reported a unimodal to transmodal gradient of functional connectivity in the former three areas, suggesting a cortical gradient of representation from sensory information to more abstract function ^26^. A similar gradient was observed during speech recognition tasks, as represented by the gradient from visual/tactile (sensory) information to emotional/social (abstract) information ^27^. In these areas, such gradients spatially correspond to our weight gradient from relatively strong emotions (e.g. ‘Sexual desire’, ‘Disgust’ and ‘Nervous scared’) to more complex emotions (e.g. ‘Awkwardness’, ‘Emotional hurt’ and ‘Interest’). This suggests that such strong emotions might be related to sensory or physical information, while more complex emotions might be related to more abstract (higher-order cognitive) information. Furthermore, in the postcentral area, we found a smooth gradient of ‘Nervous scared’ and ‘Disgust’ (including empathic pain). In this area, such localisation of these negative emotions was also reported in previous fMRI studies ^28, 29^. However, the cortical gradient of the representation has not been revealed. This area is well known to have selectivity to somatosensory information, where regions responding to upper body parts are located more ventrally ^30^. The localisation of the negative emotions may be caused by bodily reactions to the experience of high-arousal emotions, such as goose pimples ^31^, and the weight gradient may be caused by differences in the bodily parts exhibiting responses among these emotions.

In the current study, we demonstrated localisation of emotion categories in the whole cortex, but not in the sub-cortex. Previous brain-imaging studies traditionally focused on localisation of basic emotions into the sub-cortical areas ^32–36^ and the connected cortical areas such as insular and cingulate ^37–39^. In particular, the amygdala is well known to be sensitive to ‘fear’ ^40, 33^. Although our study showed emotion representation in these connected areas, we could not provide strong support for the relationships to the sub-cortical areas. For example, when we examined voxels showing high prediction accuracy of the emotion-category model, in fact such voxels were found in the amygdala, (no. of significant voxels/no. of voxels, S1:5/334; S2: 69/427; S3:11/392; S4:61/427; S5:30/398; S6:8/349; S7: 8/400; S8:5/361, uncorrected *p* < 0.0001, Pearson’s correlation test). However, the effect size was smaller than that in most cortical areas. Therefore, we here focused on the cortex.

Our weak support for association of emotion with the amygdala might have been caused by the experimental settings. In most previous studies ^41, 42, 35, 36^, brain activities were measured when evoking a specific emotion (especially ‘fear’) and also under neutral conditions, and the relationships of emotion to the amygdala were examined as a difference of activation between the two conditions. In comparison, we measured brain activities when feeling certain kinds of emotion evoked by movie scenes, and the responses to each emotion were estimated without comparison to the neutral condition. Furthermore, the amygdala activity might have been influenced by a task modality, where the difference in activation between the two conditions was lower in a movie-viewing task than in a still-image-viewing task ^36^. Therefore, we could not provide strong evidence for the relationship of emotions with the amygdala.

One important issues in this context is about the relationship between emotions and objective information. In this study, we subtracted the effect of sensory information when estimating emotion-category weights (see Methods). However, it is natural to assume that emotion is positively correlated with objective information such as visual and auditory features (e.g. ‘a baby induces feelings of adoration’, ‘a music in a minor key induces feelings of sadness’). Actually, intense competitions to predict emotions from objective information (e.g. visual, auditory, and linguistic features) are lively held in the engineering field, and the state-of-the-art methods show good prediction performance ^43–46^. To obtain a comprehensive understanding of emotion, future study is necessary to perform close comparisons of objective information, emotions, and the related brain activity.

Taken the obtained findings together, we found that a rich variety of emotion categories are represented in the human brain. In this study, we visualised the representation as a semantic space consisting of 25 emotion dimensions, each of which can be interpreted as similar emotion categories. In addition, we found smooth gradients of the emotion representation on the cortical surface, especially from unimodal to transmodal regions.

## Methods

### Subjects

Eight healthy individuals (S1–S8; age 23-32; four females) with normal or corrected-to-normal vision participated in our experiments. Before the experiments, we explained to the subjects that stimuli to be used would include extreme content, such as violent, disgusting or erotic representations. All the subjects accepted this and provided written informed consent. The ethics and safety committees of the National Institute of Information and Communications Technology approved the experimental protocol.

### Experimental design

In our experiments, fMRI BOLD responses were recorded while subjects watched audio-visual stimuli. The stimuli consisted of 138 movie clips from a video-sharing site *vimeo* (https://vimeo.com/jp), which were selected to induce a rich variety of emotions. Examples of the movie genres were as follows: horror, violent drama, comedy, romance, fantasy, daily life scenes, and action movies. Movie clips were cut down to 10– 20 s in length (mean of 15 s), and recreated as a sequence of stimuli by combining the selected clips in a random order.

The visual stimuli were presented at the centre of a projector screen with 23.3 × 13.2 degrees of visual angle at 30 Hz. The audio stimuli were presented through MR-compatible earphones with an appropriate volume level for each subject. The subjects were instructed to watch the clips naturally as if watching TV show in daily life. For each subject, fMRI data were collected in 3 separate sessions over 3 or 4 days. Each session consisted of six movie-watching runs (each run lasting 610 s). A total of 18 runs were divided into 12 model training runs and 6 model testing runs. The model training runs were used to train encoding models and consisted of 480 different movie clips shown once each (total 7,200 s). The model testing runs were used to assess model prediction accuracy and consisted of three different types of 300-s movie sequence shown four times each (total 3,600 s). None of the movies in the training runs was shown in the test runs.

### fMRI data acquisition

fMRI data were acquired using a 3T Siemens Trio TIM scanner (Siemens, Germany) with a standard Siemens 32-channel volume coil and a multiband gradient echo-planar imaging sequence ^47^ [TR = 2,000 ms, TE = 30 ms, flip angle = 62°; voxel size = 2 × 2 × 2 mm^3^, matrix size = 96 × 96, 72 axial slices, FOV = 192 × 192 mm^2^, multiband factor = 3]. Anatomical data were collected on the same 3T scanner using T1-weighted MPRAGE [TR = 2530 ms, TE = 3.26 ms, flip angle = 9°, voxel size = 1 × 1 × 1 mm^3^, matrix size = 256 × 256, 256 axial slices, FOV = 256 × 256 mm^2^].

### fMRI data preprocessing

The Statistical Parameter Mapping toolbox (SPM8, http://www.fil.ion.ucl.ac.uk/spm/software/spm8/) was used to preprocess EPI data. We performed motion correction by aligning all of the EPI data to the first image from the first scan for each subject. For each voxel, responses were normalised by subtracting the mean response across all time points. Then, long-term trends were removed by subtracting results of the median filter convolution (120-s time window). To define anatomical regions, for each subject, the cerebral cortex was segmented into 156 regions of the Destrieux atlas ^22^ by using FreeSurfer ^48^. The segmentation results in T1 space were registered to the EPI space using Freesurfer functions, and each voxel was given one anatomical label.

### Emotion ratings and preprocessing

We collected ratings regarding each of the 80 emotion categories (see ‘**80 emotion categories**’ in Methods) induced upon exposure to the movie stimuli used in our fMRI experiments. To obtain the emotion ratings, we recruited 174 annotators. They were instructed to rate how well an emotion category (e.g. ‘laughing’) matched to the movie scene, by assigning a value ranging from 0 (not match at all) to 100 (matched perfectly). These annotators were also instructed to make ratings based not on ‘movie character’s feeling’, but on ‘their own feeling’. The ratings were made by dragging a mouse while watching the movie stimuli. The ratings were stored at 1-s resolution. To obtain reliable data, first, we conducted an aptitude test for each annotator. Specifically, we used a 246-s test movie, and examined consistency of temporal fluctuation of ratings between a template by one of the authors (NK) and each annotator, regarding each of two emotions: ‘fear’ and ‘disgust’. From the results, for all the 174 annotators, the ratings showed significantly high correlations with the template ratings in both emotions (*p* < 0.05 for 246 time samples).

For each emotion category, each of four different annotators rated all of the movie stimuli. At most, two emotions were rated by one annotator, and we prevented them from rating similar emotions in successive rating periods. (In each period, a annotator rated for the whole movie stimuli.) The ratings for each annotator and each emotion were de-noised by convolving a median filter (5 s time window), and de-trended by subtracting results of convolving another median filter (150 s time window). For each emotion category, the preprocessed ratings were averaged across those for the four annotators at 2-s resolution (i.e. BOLD sampling rate). Finally, we obtained preprocessed 80-emotion ratings of 3,600 samples used as training data, and 80-emotion ratings of 1,800 samples used as test data.

### Model fitting

To estimate the BOLD-response patterns to 80 emotion categories, we constructed a voxel-wise linear regression model to explain BOLD responses ^51, 52^. The stimulus vector (total 2,080 dimensions) included emotion ratings and sensory factors (visual and auditory features). The latter was included to remove spurious correlation with the sensory factors (see **Removing spurious correlation between emotion ratings and sensory information**). To capture the hemodynamic response, the stimulus vector was concatenated with three temporal delays of 2, 4 and 6 s (total 6,240 dimensions). The model weights were optimised by least squares with L2-regularisation. The regularisation coefficient (*γ*) was optimised in 10-fold cross validation using 10 unique training-validation (9:1) subsets by randomised sampling from the training data (3,600 samples). In each cross-validation step, we then constructed the regression model using a training subset, and computed prediction accuracy using a validation subset for each *γ* of 2^*i*^, where *i* = {0,2, …, 25}. The prediction accuracy was quantified as an across-voxel average of the correlation coefficients between the actual and predicted training BOLD responses. We employed the best *γ* showing the highest accuracy across 10 repetitions. We constructed the regression model with the best *γ* using the training data (3,600 samples), and computed the prediction accuracy using the test data (1,800 samples). In our main analyses, we used voxels with high prediction accuracy of the emotion-related BOLD responses, after regressing out the sensory factors. To determine the emotion dimensions using CCA, we employed voxels with high prediction accuracies (uncorrected *p* < 0.0001) averaged across the 10 folds (3,984–13,068 cortical voxels per subject). To construct the semantic space of emotion, we employed voxels with high prediction accuracy (uncorrected *p* < 0.0001) for the test data (18,684–34,066 cortical voxels per subject).

### Removing spurious correlation between emotion ratings and sensory information

To estimate unalloyed responses to emotion categories, spurious correlation with a sensory factor was explained away from model prediction for the BOLD responses to 80 emotion categories ^52^. For this purpose, we employed low-level visual and auditory features as the sensory factor, and used them to fit the linear regression model, but these were not used in the model prediction.

As low-level visual features, we employed output of 2,139 motion energy filters ^51^. Each filter consists of quadrature pairs of spatiotemporal Gabor filters. Input frames were obtained at 15 Hz, and resized from 720 × 1280 × 3 to 96 × 171 × 3, followed by cropping in the centre to a size of 96 × 96. Then, the image was converted from RGB colour to (CIE) L*A*B* colour space, and the colour information was discarded. The motion energy signals were yielded from the filter output, and then log-transformed and averaged across 2 s (TR). Consequently, we obtained 2,139 visual features, which represent preferences to spatial frequencies, temporal frequencies, and orientations. To minimise the computational burden, we reduced the original dimensions to 1,000 using singular value decomposition. These 1,000 components preserved 83.8% of the variance explained in the original features.

As the low-level auditory features, we employed output of the modulation-transfer function model ^53^. The spectrogram was generated using 128 bandpass filters ^54^ with window size of 25 ms and hop size of 10 ms. Then, the spectrogram was convolved with quadrature pairs of modulation-selective filters for 10 spectral modulation scales and 10 temporal modulation rates. The modulation energy was calculated using the same methods as reported by Nishimoto et al. (2011) ^51^. Modulation energy was log-transformed, averaged across 2 s (TR), and further averaged within each of the 20 nonoverlapping frequency ranges logarithmically spaced in the frequency axis. From the results, we obtained 2,000 auditory features, which represent preferences to frequencies of audio signal, and the temporal variation of the preference frequencies. The same as in the visual feature extraction, we reduced the original dimensions to 1,000 to minimise the computational burden. The 1,000 components preserved 93.4% of the variance explained in the original features.

As supplementary results, we showed the prediction accuracies of the BOLD responses using each type of the three features: emotion, visual and auditory (Fig. S7). The accuracy was quantified using Pearson’s correlation coefficients between the actual and predicted responses. We confirmed high prediction accuracy for the early visual and early auditory cortex from the visual and auditory features, respectively. This indicates that these two features could plausibly explain BOLD response in the early visual and auditory cortices.

### Emotion dimensions based on the BOLD-response patterns

To estimate significant emotion dimensions, CCA was performed based on correlation in temporal fluctuation between each emotion rating and BOLD responses. In the CCA, the two types of factor loadings (***A***, ***B***) were estimated to have the maximum correlation between the linear combinations of emotion ratings and the BOLD responses. Using the training data, we estimated the factor loadings (***A^∗^***, ***B***^∗^) as follows:

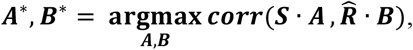

where ***corr***(***i***, ***j***) denotes correlation coefficients between ***i*** and ***j***. ***S*** (3,600 × 80) is the emotion ratings. ***R̂*** (3,600 × 1,800) is the dimension-reduced BOLD responses of voxels showing good prediction accuracy (correlation coefficients, *p* < 0.0001) in 10-fold cross-validation in the linear regression model (see ‘A linear regression model for 80 emotion categories’ in Methods). To validate the estimated ***A***^∗^ and ***B***^∗^, we used test ***R***̂ (1,800 × 1,800) and ***S*** (1,800 × 80). Then, we quantified the significance of each dimension of ***A*** as a Pearson’s correlation coefficient between each dimension of ***S*** · ***A*** and that of ***R***̂ · ***B***. The significance was defined by the statistical significance (*p* < 0.01; with Bonferroni correction for 80 emotion categories). After obtaining significant dimensions of ***A***^∗^ (***A_sig_***), we conducted a varimax factor rotation ^19^ on ***A_sig_*** to explain dimensions with fewer emotion categories. The ***A_sig_*** provided interpretation of a specific BOLD-response pattern by using the correlated emotion categories. Then, we calculated the median of the numbers of the significant dimensions across subjects (25 dimensions, Fig. S1). The number (25) was used in the subsequent analysis to determine the number of dimensions for a semantic space of emotion.

### A semantic space for brain representation of 80 emotion categories

To construct a semantic space, we used emotion-category weights of the regression model using voxels with high prediction accuracies (*p* < 0.0001, uncorrected; the number of significant voxels ranged from 18,684 to 34,066 cortical voxels for each subject). The emotion-category weights were averaged across three temporal delays, and we obtained weight matrices (18,684 –34,066 cortical voxels x 80 emotions) that represent voxel-wise selectivity to each of the semotion categories. The weights for each subject were concatenated across eight subjects. We called them ‘group weights’. PCA was used to reduce the dimensions of the group weights to the number of significant dimensions of the CCA results (25 dimensions). We defined the PCA space as the semantic space of emotion. Then, we conducted a varimax factor rotation on the principal components, to obtain interpretation of each emotion data of dimension. Factors with high negative loadings were rotated to have the opposite direction by multiplying by −1. The rotated components were used as ‘25 emotion dimensions’ in analyses for the cortical gradient of emotion representation.

To confirm across-subject consistency of a semantic space, we performed a leave-one-subject-out method. For this, we constructed a semantic space for a single subject (individual space), and also that for the weights concatenated across the remaining seven subjects (sub-group space). As a control semantic space, we also constructed a semantic space by performing PCA (category-dimension reduction) on the emotion ratings of the training data. All of the three types of semantic space consist of the number of significant dimensions in the CCA analysis for the left-out subject (19–32 dimensions per subject). To support across-subject consistency of individual spaces, we showed that the similarity to the sub-group space (*r_g_*) was higher than that to the control space (*r_c_*) for each subject. To quantify the similarity between semantic spaces, we calculated a Pearson’s correlation coefficient in emotion distributions between each semantic space pair. The emotion distribution was quantified as pair-wise distances (correlation distance) of emotion categories in each semantic space.

### Cortical gradients of emotion dimensions

To visualise how emotion-category weights were distributed through the cortical surface, we used the first three components of the semantic space. Emotion-category weights of each voxel were first projected to the semantic space consisting of the three components. Then, the projected coordinates were normalised to range from 0 to 1 by z-scoring and linear scaling. The output result was used as RGB colour projected to voxel coordinates in the flattened cortical map. We can observe the weight gradient of emotion categories as the RGB colour gradient on the cortical map. Furthermore, we assigned an RGB colour to each emotion category by using three normalised components for each emotion category. This colour was used for visualisation of the semantic space in Fig. 2a. Each of the emotion-category colours implies an association with regions which have a similar colour in the flattened cortical map.

In the cortical map for emotion-category weights, we observed a smooth gradient in four areas: the postcentral area, the superior temporal area, the inferior parietal area, and the precuneus. To quantify the weight gradient, for each voxel, we first computed the weight of the 25 emotion dimensions by multiplying the 80 dimensional weights [1 × 80] and the 25 emotion dimensions [80 × 25]. Then, we manually defined eight lines on a smooth gradient in the four areas at each of the left and the right hemispheres (see Fig. 4a), with reference to the flattened cortical map using the in-house Matlab (MathWorks Inc.) GUI toolbox. To confirm that each line is located at similar anatomical location, we obtained the anatomical label of the Destrieux atlas ^22^_-_from each line for each subject (Fig. S4). The label was defined based on anatomical location using Freesurfer ^48^.

### Eighty emotion categories

The 80 emotion categories are listed below:

(1)love, (2)amusement, (3)craving, (4)joy, (5)nostalgia, (6)boredom, (7)calmness, (8)relief, (9)romance, (10)sadness, (11)admiration, (12)aesthetic appreciation, (13)awe, (14)confusion, (15)entrancement, (16)interest, (17)satisfaction, (18)excitement, (19)sexual desire, (20)surprise, (21)nervousness, (22)tension, (23)anger, (24)anxiety, (25)awkwardness, (26)disgust, (27)empathic pain, (28)fear, (29)horror (bloodcurdling), (30)laughing, (31)happiness, (32)friendliness, (33)ridiculousness, (34)affection, (35)liking, (36)shedding tears, (37)emotional hurt, (38)sympathy, (39)lethargy, (40)empathy, (41)compassion, (42)curiousness, (43)unrest, (44)exuberance, (45)appreciation of beauty, (46)fever, (47)scare (feel a cill), (48)daze, (49)positive-expectation, (50)throb, (51)sexiness, (52)indecency, (53)embarrassment, (54)oddness, (55)contempt, (56)alertness, (57)eeriness, (58)positive-emotion, (59)vigor, (60)longing, (61)tenderness, (62)pensiveness, (63)melancholy, (64)relaxedness, (65)acceptance, (66)unease, (67)negative-emotion, (68)hostility, (69)levity, (70)protectiveness, (71)elation, (72)coolness, (73)cuteness, (74)attachment, (75)encouragement, (76)annoyance, (77)positive-fear, (78)aggressiveness, (79)distress, (80)stress

## Acknowledgements

This study was partly funded by Brother Industries Ltd. and JSPS KAKENHI Grant Numbers 15H05311, 15H05710 and JP18H05091 in #4903 (Evolinguistics). Data were collected with support from the National Institute of Information and Communications Technology.

## Author contributions

N.K. and S.N. designed the experiment; N.K collected the data; N.K analysed the data with support from T.N. and S.N. and all authors wrote the manuscript.

## Data availability

The data that support the findings of this study are available from the corresponding author upon request.

